# The impact of C-Tactile Low threshold mechanoreceptors on affective touch and social interactions in mice

**DOI:** 10.1101/2021.01.13.426492

**Authors:** Emmanuel Bourinet, Miquel Martin, Damien Huzard, Freddy Jeanneteau, Pierre-Francois Mery, Amaury François

## Abstract

Affective touch is necessary for proper neurodevelopment and sociability. However, it is still unclear how the neurons innervating the skin detect affective and social behaviours. To clarify this matter, we targeted a specific population of somatosensory neurons in mice, named C-low threshold mechanoreceptors (C-LTMRs), that appears particularly well suited physiologically and anatomically to perceive affective and social touch but whose contribution to these processes has not yet been resolved. Our observations revealed that C-LTMRs functional deficiency from birth induced social isolation and reduced tactile interactions in adults. Conversely, transient increase in C-LTMRs excitability in adults using chemogenetics was rewarding, temporally promoted touch seeking behaviours and thus had pro-social effects on group dynamics. This work provides the first empirical evidence that specific peripheral inputs alone can drive complex social behaviour, demonstrating the existence of a specialised neuronal circuit originating from the skin wired to promote interaction with other individuals.

## Introduction

The rewarding value that emerges from touch is essential for decision making and motivation, especially in social animals, and dysregulation of this process leads to debilitating psychiatric or neurologic conditions, including autism, anxiety or depression ^1–3^. Nonetheless, the neural mechanisms underlying emotional tactile sensory processing relationship with social behaviour are at the early stage of their understanding.

The skin is innervated by an array of functionally distinct populations of receptors contributing to touch, which can be distinguished by their response properties, activation threshold, conduction velocity, and the type of end organ that they innervate ^4^.

One class of these receptors, the C-Tactile fibers (CT), are particularly well responsive to tactile stimulation categorized as “pleasant” and “affective”, by human subjects ^5,6^. These neurons are unmyelinated low-threshold mechanoreceptors (LTMRs) and respond to non-noxious touch with a predilection for slow-moving and low-force, stroking stimuli, such as gentle brushing ^6–8^. Activation of CTs in humans provides poor conscious spatial and qualitative information to the subjects, who nonetheless still carry a positive feeling related to the gentle brushing of the skin ^9^. Furthermore, the positively valent tactile information conveyed by CTs make them particularly well suited to link tactile information to social bonding ^10,11^.

Evidence for pleasant touch contribution to sociability and related disorders was also found in laboratory animal models. For example, playful and pleasant touch is rewarding and contributes greatly to social development in adolescent rats ^12^. In mice, two recent studies linked the disruption of LTMR functions with alterations of social behaviour typical of autism spectrum disorder (ASD) ^13,14^. Specifically, these studies used genetically modified mice recapitulating mutations found in human patients within the *Mecp2, Shank3B*, and *Fmr1* genes. Peripherally restricted conditional Knock-Out (KO) of these genes greatly altered Low-threshold-mechanoreceptors (LTMRs) function which induced multiple phenotypes associated with ASD, especially social deficits that are usually observed in constitutive KOs of these genes. However, LTMR sensory neurons are known to be a highly heterogeneous population, leaving open the question regarding the contribution of CT and pleasant touch to these social deficits.

We and others genetically identified a population of primary sensory neurons in mice, named C-Low-threshold-mechanoreceptors (C-LTMRs), with similar functional properties than CTs. These studies characterized genes specific to C-LTMRs in rodents such as TAFA4, but also a set of genes allowing to differentiate C-LTMRs from other sensory neurons, such as tyrosine hydroxylase (TH), VGlut3, TAFA4 or Ca_v_3.2, that may be used to gain genetic access to these neurons ^15–20^. These studies unveiled the contribution of C-LTMRs, and these genes, to pain chronification in the context of neuropathic or inflammatory pathological pain. However, none of these studies considered the role of C-LTMRs in touch sensation, within naturalistic conditions and especially their role in social behaviours.

In the present study, we explored the role of C-LTMRs in social interactions. Using two mouse transgenic models to decrease or facilitate C-LTMRs excitability, and a new tracking system to automatically annotate social behaviour in groups of 4 animals, we clarified the specific function of C-LTMRs in rodent inter-individual relations.

## Results

### Social behaviours are impaired in Cav3.2^Nav1.8^cKO

First, we aimed at investigating the consequence of C-LTMRs hypofunction on social behaviour. For that purpose, we used a genetic mouse model where the expression of the low threshold calcium channel Ca_v_3.2 is conditionally knocked out in C-LTMRs, by crossing Ca_v_3.2 ^GFP-flox^KI with Na_v_1.8^cre^ mice as we previously described ^16^. In this mouse model (Cav3.2^Nav1.8^cKO), C-LTMR impaired function starts just before birth. Indeed, removing Ca_v_3.2 expression from C-LTMRs increases the firing threshold of action potentials, reduces the firing frequency and lowers the conduction velocity of these fibres transforming their mechanical sensitivity into High Threshold Mechanoreceptors (HTMR) phenotype ^16^.

To evaluate the consequences of C-LTMRs deficiency on social preference behaviour, we used the three-chamber paradigm (ie. Crawley test). Interestingly, when compared to control Ca_v_3.2^GFP-flox^KI littermates, male Cav3.2^Nav1.8^cKO mice spent less time interacting with an unfamiliar mouse than with an unanimated object, which is summarized by the reduction in the preference index (**Figure 1a, Supplementary Fig. 1a**). To further investigate the precise quality of the social and tactile interactions that appears to be impaired in this model we used a novel paradigm where mice could interact freely with each other. The live mouse tracker (LMT), based on a machine learning analysis framework, was designed by de Chaumont and colleagues for that specific purpose ^21^. This system allows the tracking and automatic annotation of mice behaviours and social interactions in their environment for multiple days ^21^. Using this system, we analysed the behaviour of 5 groups of 4 male mice (each group composed of 2 controls Ca_v_3.2^GFP-flox^KI, and 2 C-LTMRs-impaired Cav3.2^Nav1.8^cKO, all littermates) for three consecutive nights (**Figure 1b**). For every time frame, the LMT detects head, tail, ears, eyes and nose position, producing a geometrical mask for each mouse. These data permit the computation of different behavioural events based on mice geometries movements and localization compared to other mice. Overall, we detected 28 events that were separated into seven categories: body configuration, isolated behaviour, position in contact, type of contact, social configuration, social approach and social escape. These events were analysed separately for both their number (**Figure 1c**) and their duration (**Figure 1d**) for each individual. To compare C-LTMR-impaired mice and their control cagemates with the same baseline and to reduce inter-experiment variability, the value of each behavioural trait for one Cav3.2^Nav1.8^cKO mouse was compared with the mean level of this trait from the two control cagemates (called LMT index thereafter). An LMT index value above one indicates that the Cav3.2^Nav1.8^cKO mice perform more occurrences of a specific trait compared to controls whereas a value below one means that the mutant mice do less of that trait than controls.

**Figure 1:**
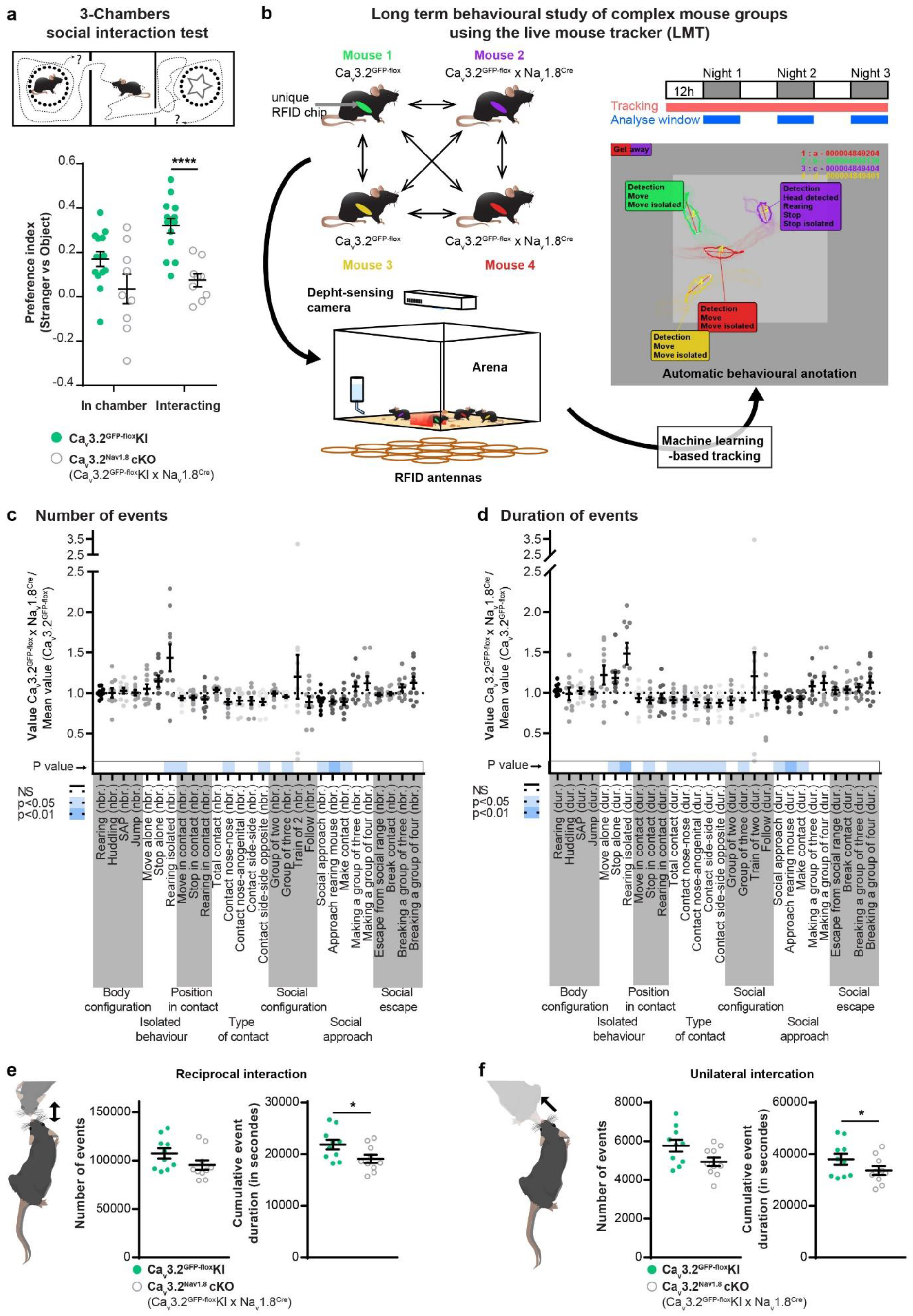
C-LTMRs impairment via Ca_v_3.2 deletion reduces social behaviour. a. In a three-chamber social interaction test, control Ca_v_3.2^GFP-flox^KI littermates have a significantly higher preference index toward a stranger mouse than Cav3.2^Nav1.8^cKO. Unpaired t-Test p<0.001, n Cav3.2^GFP-flox^KI= 14 (light green circles), n Cav3.2^Nav1.8^cKO = 9(open grey circles). b. Live mouse tracker configuration description. Each animal is implanted with a RFID chip for authentication and tracking. The system includes 16 RFID antennas that cover the 50 by 50 cm arena which allow, with the deep sensing camera and the machine learning algorithm, to track and automatically annotated the behaviour of 4 mice. Each group of mice includes 2 Ca_v_3.2^GFP-flox^KI and 2 Cav3.2^Nav1.8^cKO. the experiment was performed over 3 days and each behaviour was analysed during the nocturnal activity phases. c. LMT index for 28 behavioural traits. Sum of all the number of events for the three nights. d. LMT index for 28 behavioural traits. Sum of the duration of events for the three nights. For c. and d.: The index for each trait was compared to one using one-sample two-sided Student’s t-tests (corrected for multiple testing, because 28 tests were conducted for each strain). The p values were color-coded in shades of blue depending of the p value as indicated on the left. n = 10 e. f. Raw values extracted from the LMT for three consecutive nights for Ca_v_3.2^GFP-flox^KI (light green circles; n =10) and Cav3.2^Nav1.8^cKO (open grey circles; n=10). e. number of events and cumulative duration of reciprocal social interaction (∗ p=0.0372). f. number of events and cumulative duration of unilateral social interaction (∗ p=0.0429). unpaired t-test

The LMT-index indicates that Cav3.2^Nav1.8^cKO mice spent more time isolated than controls (time *stop alone*: +18.2%±7), **Figure 1d**), without any noticeable differences in locomotor activity (**Supplementary Fig. 1b**) or exploratory behaviour (stretch attending posture: (*SAP*) **Figure 1e&d**). In social events, C-LTMR impaired mice showed a small, but statistically significant, decrease in time spent engaged in all type of contacts (−15.7%±3.2 in average per interaction bout, **Supplementary Fig. 1c**). Moreover, the duration of reciprocal (here associated with nose-to-nose contacts and side-by-side contacts) and unilateral social interactions (associated in these experiments with giving ano-genital contacts) were reduced, even if the number of events is not statistically different (**Figure 1e&1f**). However passive social interaction (associated in our experiment with receiving ano-genital contacts) were not altered (**Supplementary Fig. 1d**). Cav3.2^Nav1.8^cKO mice also displayed shorter social approaches leading to a contact (*social approach*: −10.9%±1.9 and *make contact*: −7.3%±2.8, **Figure 1d**) and spent less time in groups of three mice (−9.27%±1.9) (**Figure 1d**). However, no difference was observed in the social escape behaviours category. Taking together, the results from the 3-chamber test and the LMT revealed a deficit in sociability of Cav3.2^Nav1.8^cKO mice compared to their control littermates.

### A new viral strategy to specifically target C-LTMRs in mice

Next, we deepened our investigations on C-LTMR role in sociability by designing a chemogenetic strategy to selectively excite C-LTMRs remotely in adult mice independently of Ca_v_3.2 protein function and, importantly, without any postnatal functional perturbation of C-LTMR. Our strategy to target C-LTMRs in adult mice, illustrated in **Figure 2a and b**, consists on expressing a gene of interest under the control of the mini-Ca_v_3.2 promoter in a Cre-dependent manner. Indeed, C-LTMRs can be defined by the expression of both the sodium channel Na_v_1.8 and the calcium channel Ca_v_3.2 ^16,22^. We previously engineered adeno associated viruses (AAV) with a mini-Cav3.2 sequence and validated its faithful expression in Cav3.2 positive neurons within the dorsal horn of the spinal cord ^23^. Here we used AAV-PHP-S serotype that has a high tropism for peripheral sensory neurons ^24^, which we delivered into Nav1.8^Cre^ heterozygote mice. This intersectional strategy combining the expression pattern of Cav3.2 and Nav1.8 aims at restricting the expression of the viral payload into C-LTMRs. To achieve C-LTMRs chemogenetic stimulation with this strategy, we inserted within the pAAV vector the HA tagged-hM3Dq excitatory DREADD cassette.

**Figure 2:**
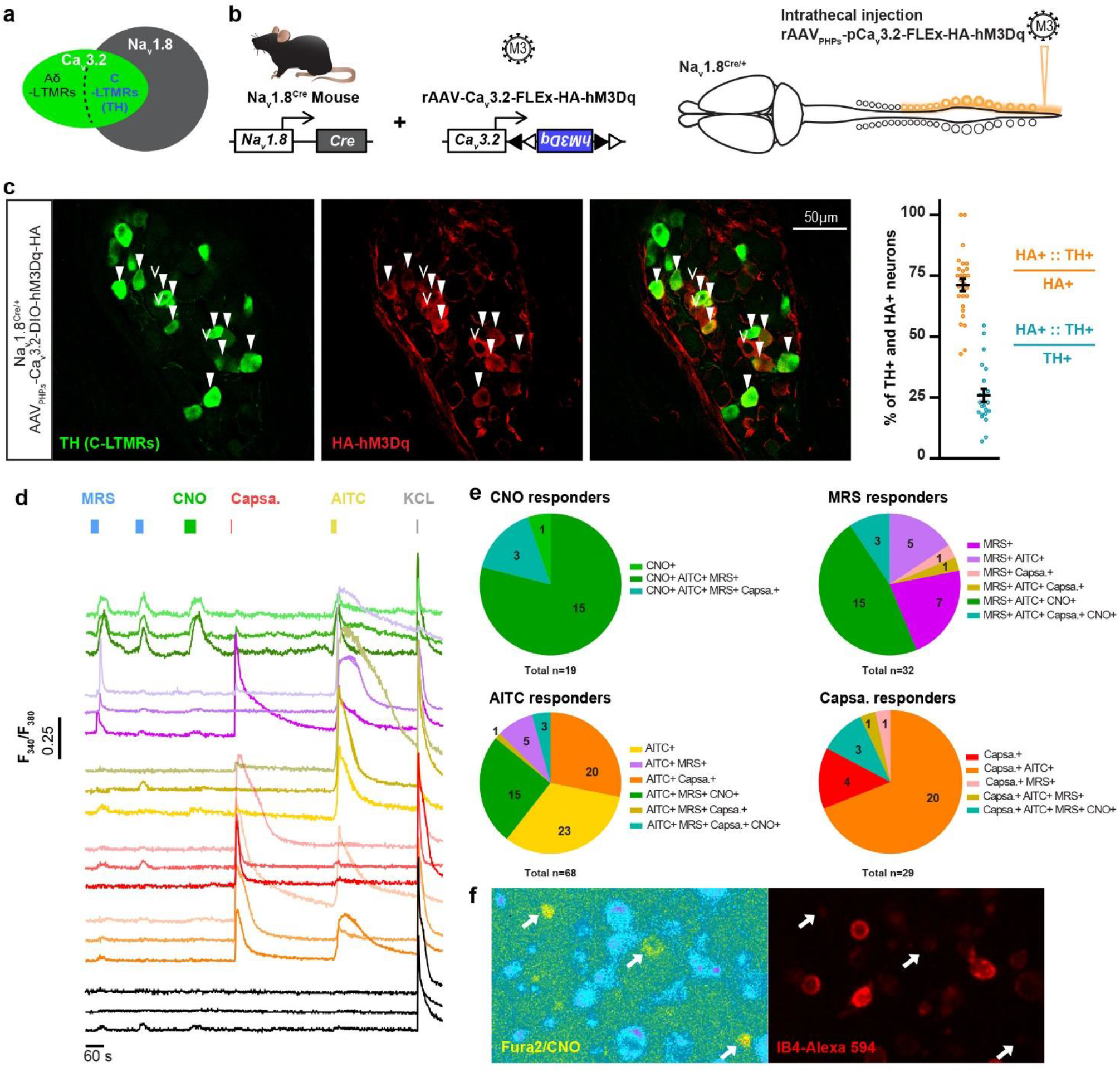
Viral expression of DREADD receptors HA-hM3Dq in C-LTMRs. a. Diagrams representing primary sensory neurons expressing Na_v_1.8 and Ca_v_3.2 and the population defined by the expression of both ion channels. b. Strategy to express HA-hM3Dq in C-LTMRs using intrathecal injection of AAV_PHPs_ serotypes in Nav1.8^Cre^ mice. c. Left: representative images of an Immunofluorescence of Tyrosine Hydroxylase (TH, C-LTMRs marker, green) and HA tag (HA-hM3Dq, red) in thoracic dorsal root ganglia (T13) of a Na_v_1.8Cre mice injected with the AAV_PHPs_-pCa_v_3.2-FLEx-HA-hM3Dq. Filled arrowheads indicate examples of neurons considered positive for HA and TH. Empty arrowheads indicate examples of neurons considered positive for HA only. Right: Bar graph of the percentage of neurons positive for HA and TH immunostaining over the all HA positive population (Orange dots) or over the all TH positive population (Tile dots) in thoracic and lumbar ganglions. N=3 mice; Thoracic: T7 to T13; Lumbar: L1 to L6. d. Representative individual calcium influx in cultured DRG neurons using Fura2 following bath perfusion of MRS2365 (200nM, blue) CNO (30µM, green), Capsaicin (Capsa., 500nM, red) AITC (200µM, yellow) or KCL (40mM, Grey). e. Pie charts of the number of neurons responding to the different chemical compounds. Based on the amplitude of the calcium influx responses, neurons were classified as CNO responder (Top Left), MRS responder (Top right), AITC responder (bottom left) and Capsaicin responders). A total of 168 neurons were recorded from 3 animals. f. representative images of ratiometric calcium imaging on cultured DRG from a Na_v_1.8Cre mice injected with the AAV_PHPs_-pCa_v_3.2-FLEx-HA-hM3Dq. Left panel: ratiometric Fura2 after CNO bath perfusion, right panel: live IB4-Alexa 594 sainting of the same field of view. Arrows indicates CNO-responsive DRG neurons.

HEK293T cells were transfected with the pAAV vector (pAAV-pCav3.2-FLEx-HA-hM3Dq) to validate its Cre-dependency and functionality. As the Cav3.2 promoter activity is highly enhanced by the EGR1 transcription factor ^25^, we co-expressed the murine EGR1 cDNA and added or not the Cre recombinase (Cre-GFP fusion). We analysed the DREADD functionality using Calcium fluorimetry in Fura2 loaded Cre-GFP positive cells. In a representative field, out of 102 cells recorded, 28 were GFP positive and of those 17 responded to the hM3Dq pharmacological actuator Clozapine-N-oxyde (CNO) (**Supplementary Fig. 2a & b**). No responses to CNO were observed in non-GFP cells.

We then packaged viral particles with this validated construct using the AAV-PHP-S capsid. Although AAV PHP-S can be delivered systematically to access sensory neurons, we locally injected the virus intrathecally with minimal invasiveness and a sole access to DRGs and no other sensory neurons such as those of the vagal ganglia (**Figure 2b**)^24^. Consistent with an efficient C-LTMR targeting strategy, intrathecal injection of this viral construct leads to the expression of the excitatory DREADD receptor HA-hM3Dq in C-LTMRs labelled by the tyrosine hydroxylase (TH), 8 weeks post injection (**Figure 2c**). Overall, 71.2%±2.4 of HA-hM3Dq cells were also positive to TH (N=5 mice). Inversely, HA-hM3Dq is present in 25.9%±2.6 of TH positive dorsal root ganglia (DRG, T7 to L6) neurons (**Figure 2c**). In addition, no observation was made of any specific HA immunostaining in the dorsal horn of the spinal cord (**Supplementary Fig. 2c**)

To further confirm that hM3Dq was selectively expressed in C-LTMRs accordingly to the defined strategy and kept its pro-excitatory nature, we evaluated the effect of CNO on cultured DRG neurons from animal injected intrathecally with the rAAV_PHPs_-pCa_v_3.2-FLEx-HA-hM3Dq cells were loaded with the Fura2 radiometric calcium indicator, and labelled with red-dye conjugated IB4 to access the pharmacological and functional properties of large population of neurons at the same time. Application of CNO (30µM) induced intracellular calcium increase in neurons also responding to the TRPA1 agonist, allyl isothiocyanate (AITC, 200µM), and to the P2Y1R agonist, MRS2365 (200nM) (**Figure2d and 2e**). Of all CNO responsive neurons, the large majority responded to AITC and MRS (15), 3 also responded to Capsaicin on top of AITC and MRS, and only one was not responding to anything else. Among the MRS responders, 56.2% were also responding to CNO while among AITC responders only 26.4% were CNO responders. In mouse DRGs, TRPA1 is weakly expressed in C-LTMRs and P2YR1 receptor is only expressed in C-LTMRs and TrkB positive A*δ*-LTMRs ^18,20^. As TRPA1 is not express in TrkB neurons, responses to both MRS and AITC can only be observed in C-LTMRs. Moreover, none of the CNO responsive neurons were labelled by IB4 (**Figure 2f**), in agreement with the lack of reactivity of the lectin in mouse TH positive C-LTMR neurons ^26^. Taken together, our morphological and functional data provide supporting evidence toward a specific expression of HA-hM3Dq in a large population of C-LTMRs (10.7% of all DRG neurons recorded, N= 3 mice). Accordingly, we used this experimental approach in vivo to investigate the impact of C-LTMRs stimulation in social behaviours.

### Effect of exogenous C-LTMR activation on somatosensory perception

First, we assessed the consequences of C-LTMRs exogenous activation on somatosensory perception. Eight weeks after intrathecal injections of rAAV_PHPs_-pCa_v_3.2-FLEx-HA-hM3Dq (named further: C-LMTRs^hM3Dq^) or rAAV_PHPs_-CAG-mCherry (Control) in Na_v_1.8^cre^ mice, CNO was administrated intraperitoneally (1mg/kg, IP) to all animals. 30 to 45 minutes after, a decrease in reflex paw withdrawal frequency to low force von Frey stimulation (0.07g) was observed, but not for higher forces (0.6g and 2g) or brushing in C-LMTRs^hM3Dq^ mice when compared to control mice (**Supplementary Fig. 3a**). To note, CNO injection did not trigger any alteration of motor activity or spontaneous nocifensive behaviours, such as paw shaking, guarding, grooming, licking or jumping compared to the control group.

As C-LTMRs have been implicated into temperature perception ^15,16,27^, we probed thermal sensitivity of control and C-LMTRs^hM3Dq^ mice in the thermal gradient test. 30 minutes after CNO injection, mice were placed into a 1.5m corridor with the floor at one extremity cooled down to 5°C and the other heated up to 50°C, creating a gradient of temperature between the two extremities. Once placed in the corridor, mice were allowed to explore the thermal gradient to reveal their thermotaxis behaviour. The exploration was tracked during 90 minutes and the animal’s position was annotated according to the temperature zones they visited (**Figure 3a**). When compared to control animals, CNO treated C-LMTRs^hM3Dq^ mice settled more quickly at the comfort temperature of 30°C (**Figure 3c and 3d**) and spend more time overall at this temperature (**Figure 3b and 3f**), without affecting locomotor (**Supplementary Fig. 3b and 3c**) nor the preferred temperature that was similar between the two groups (**Supplementary Fig. 3d**). This interesting result suggests that exogenous activation of C-LTMRs can reinforce motivational behaviour towards a pleasurable somatosensory stimulation.

**Figure 3:**
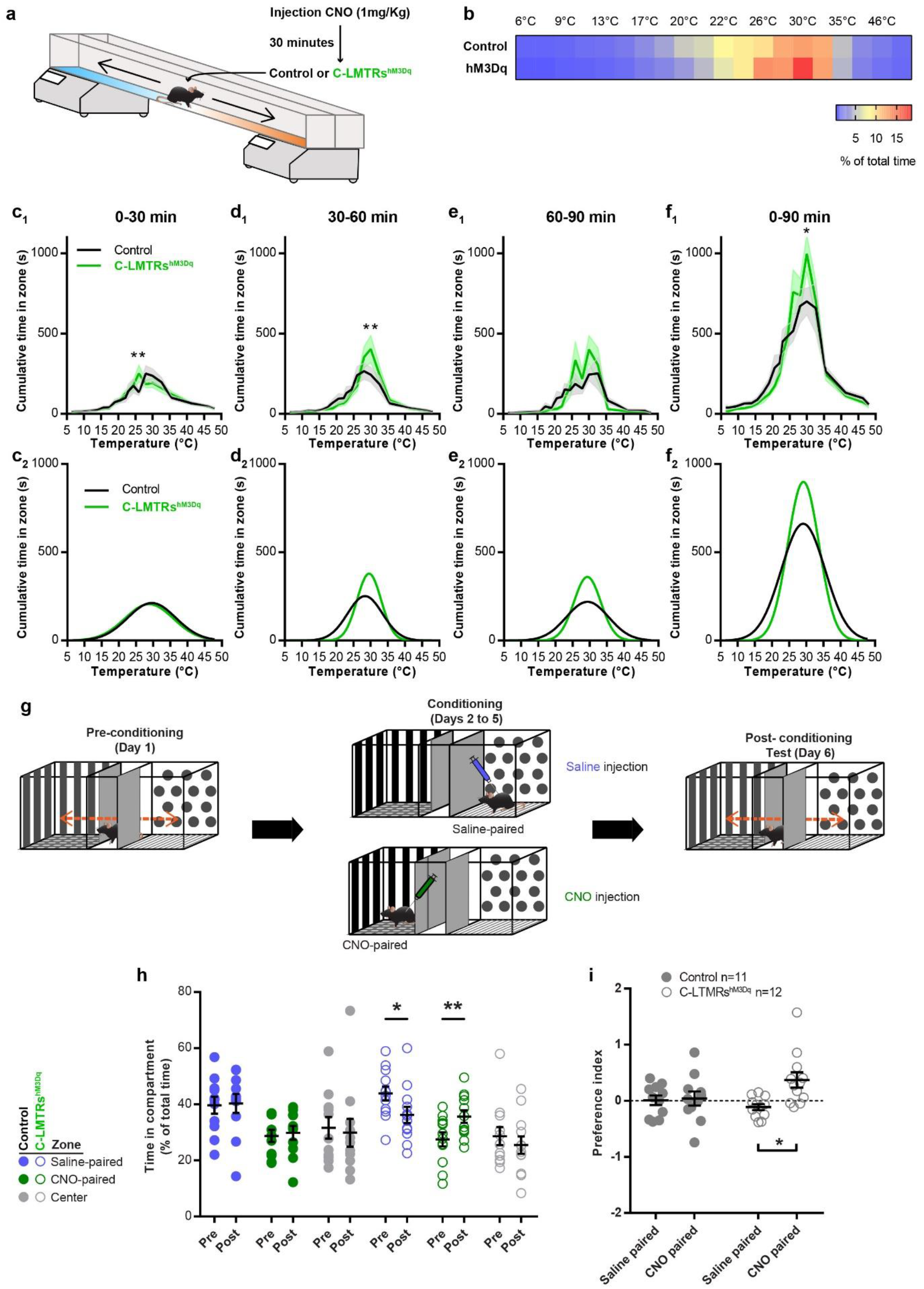
C-LTMRs exogenous activation increases thermal preference, and induced conditioned place preference. a. Thermal gradient protocol to assess thermotaxis of Control (Nav1.8^Cre^; AAV_PHPs_-CAG-mCherry; in black) and C-LTMRs^hM3Dq^ (Nav1.8^Cre^; AAV_PHPs_-pCa_v_3.2-FLEx-HA-hM3Dq; in green) mice. b. Heatmap of the zone occupancy (in % or total time) in the thermal gradient for 90 minutes. c1. Cumulative time in the thermal gradient zones during the first 30 minutes. c2. Predictive curves based on a Gaussian fit of the results in c1. d1. Cumulative time in the thermal gradient zones between 30 and 60 minutes. d2. Predictive curves based on a Gaussian fit of the results in d1. e1. Cumulative time in the thermal gradient zones between 60 and 90 minutes. e2. Predictive curves based on a Gaussian fit of the results in e1. f1. Cumulative time in the thermal gradient zones for the all 90 minutes. f2. Predictive curves based on a Gaussian fit of the results in f1. (two-way ANOVA, Bonferroni post hoc test, ∗∗p = 0.0037; ∗∗p=0.0019; ∗p=0.0112; Control n=11, C-LTMRs^hM3Dq^ n =12). g. Conditioned placed preference protocol. h. Time spent in compartments (in % of total time) before (pre-) and after (post-) 3 days of conditioning. Wilcoxon matched-pairs signed rank test (∗∗p = 0.0093; ∗p=0.0210; Control n =11, C-LTMRs^hM3Dq^ n =12). i. Conditioned placed preference index (time post - time pre/time pre). (two-way RM ANOVA, Bonferroni post hoc test, ∗p = 0.0141)

### Positive valent information is associated with C-LTMRs stimulation

As activation of C-LTMRs may results in positive feeling, next we investigated whether C-LTMRs activation could be rewarding on its own by using the conditioned place preference paradigm (CPP). Following one day of habituation to the CPP arena, control and C-LMTRs^hM3Dq^ mice were conditioned to receive saline injection (CNO vehicle) in the compartment they preferred during habituation and CNO (1mg/kg in saline) in the other compartment, for 3 consecutive days (**Figure 3g**). On the last day, mice were free to explore the entire arena and their position was video tracked. While control animals did not develop a preference for the side in which they either received saline or CNO injections, C-LMTRs^hM3Dq^ mice showed a marked preference for the compartment associated with CNO injections (32%±1.4 increase, **Figure 3h and 3i**). Overall, these two experiments suggest that activation of C-LTMRs is rewarding and can increase the rewarding value of other sensory modalities.

### C-LTMR stimulation induces touch seeking and pro-social behaviours

Taking into consideration the intrinsic emotional value conveyed by C-LTMRs activation, we finally investigated whether C-LTMR activation can affect social behaviours and social group organization. The LMT system was again used to analyse the behaviour of 5 groups of 4 mice independently during 3 nights. Each group was composed of 2 Na_v_1.8^cre^ mice injected with rAAV_PHPs_-CAG-mCherry (Control) and 2 mice injected with rAAV_PHPs_-pCa_v_3.2-FLEx-HA-hM3Dq (C-LTMRs^hM3Dq^), all male littermates. The first 24 hours were used as a habituation phase, and just before the second dark cycle, we injected either a saline (CNO vehicle) or CNO solution (IP, 1mg/kg in saline) to all animals (Control and C-LTMRs^hM3Dq^). At the beginning of the third dark cycle, groups that received a saline solution for the second dark cycle received CNO and *vice versa* (**Figure 4a**). As for Cav3.2^Nav1.8^cKO mice, we calculated the LMT-index by taking the value of each trait for one C-LMTRs^hM3Dq^ mouse and compared it with the mean level of that trait for the two control cagemates (**Figure 4 f to i**). The LMT-index was calculated from cumulated value for both saline and CNO injection during the first half of the dark cycle, at 30 minutes, 60 minutes, 90 minutes, 150 minutes (2.5 hours), 210 minutes (3.5 hours), 270 minutes (4.5 hours), 330 minutes (5.5 hours), and 390 minutes (6.5 hours) post injection. In addition, the LMT-index was also calculated at 1200 minutes during the following light cycle (20 hours) post injection. To highlight the behaviours that were exacerbated or inhibited by activation of the C-LTMRs following CNO injection, we then computed another ratio for each behaviour by subtracting the LMT-index^CNO^ to LMT-index^saline^. The p-value of these ratios was then calculated for each different time points and represented as a two-colour gradient heatmap (**Figure 4b, c d and e**).

**Figure 4:**
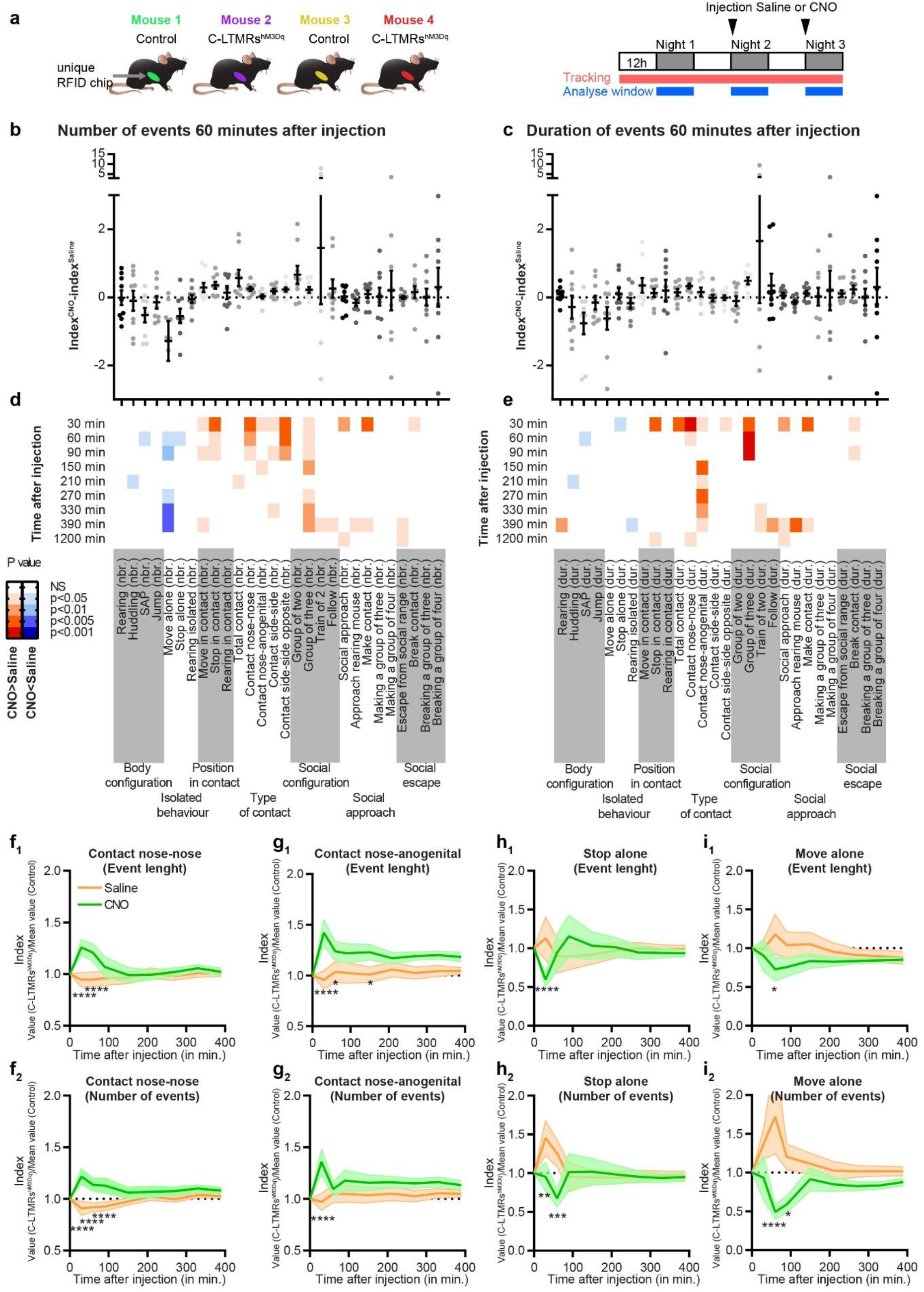
C-LTMRs exogenous activation transiently increases contact between animals and reduces isolated behaviour. a. Left: Cage composition of each group analysed in the LMT: 2 Na_v_1.8^cre^ mice injected with rAAV_PHPs_-CAG-mCherry (Control) and 2 mice injected with rAAV_PHPs_-Ca_v_3.2-FLEx-HA-hM3Dq (C-LMTRs^hM3Dq^). Right: Animals were recorded for 3 consecutive days, and received on injection of saline at the beginning of the second night cycle and one injection of CNO (1mg/kg) at the beginning of the third, or vice versa. b. LMT index^CNO^-LMT index^saline^ ratio of the number of events 60 minutes after injection for each behavioural trait automatically annotated by the LMT. c. LMT index^CNO^-LMT index^saline^ ratio of the events duration 60 minutes after injection for each behavioural trait automatically annotated by the LMT. d. and e. P values obtained by comparing the index for each trait to zero, at 9 different time points post injection (30 minutes, 60 minutes, 90 minutes, 150 minutes (2.5 hours), 210 minutes (3.5 hours), 270 minutes (4.5 hours), 330 minutes (5.5 hours), 390 minutes (6.5 hours) and 1200 minutes (20 hours) using one-sample two-sided Student’s t-tests (corrected for multiple testing). The P values were color-coded as indicated on the left depending of the level of significativity and the polarity of the mean used for comparison: negative (shade of blue) or positive (shade of red). n = 10. f to i. LMT index obtained at 30, 60, 90, 150, 210, 270, 330, and 390 minutes post saline injection (orange) or post CNO injection (green) for the cumulative duration (f_1_) and the cumulative number (f_2_) of contact nose to nose, the cumulative duration (g_1_) and the cumulative number (g_2_) of contact nose anogenital and, the cumulative duration (h_1_) and the cumulative number (h_2_) of events stop alone and their length, or the duration (i_1_) and the number (i_2_) of event move alone. two-way RM ANOVA, Bonferroni post hoc test, ∗p < 0.05 ; ∗∗ p<0.01; ∗∗∗ p<0.005; ∗∗∗∗p<0.0001

The index ratios indicate that, overall, CNO but not saline injection, significantly reduced the number of isolated events while it increased inter-individual events in C-LMTRs^hM3Dq^ mice and not in controls 1 hour post-injection (*Move alone*: −122%±2.1, 60 minutes post CNO injection; *stop alone*: −57%±1.2, 60 minutes post CNO injection; *move in contact*: +24%±0.7, 60 minutes post CNO injection; *stop in contact*: +30%±0.7, 60 minutes post CNO injection; **Figure 4b, c, d, e, h and I; Supplementary Fig. 4 d and e**). It is noteworthy that a single injection of CNO was sufficient to significantly decrease the number of events corresponding to isolated movement up to 6.5 hours following the injection (**Figure 4b and i**).

C-LMTRs^hM3Dq^ mice also appeared to be engaging more and, for longer periods, in all the different types of contact post CNO injection. We also observed that some behavioural traits were transiently altered, lasting for up to 30 to 90 minutes, while other were more durable (up to 6.5 hours post injection, a time point largely exciding the CNO clearance ^28^). Specifically, behavioural traits associated with the time spent on social exploration were significantly increased immediately after CNO injection and lasted up to 6 hours (duration of contact *nose-anogenital*: +20.4%±0.7 at 60 minutes post CNO, number of *group of three*: +22% at 60 minutes post CNO; **Figure 4d, e and g**), whereas other behaviours were only significantly increased for the first 60 to 90 minutes post CNO injection (duration and number of *contact nose-nose*: +25%±0.4 and +20%±0.3 respectively 60 minutes post CNO injection, contact *side-by-side*: +22.2%±0.8 90 minutes post CNO injection, and contact *side-by-side opposite*:+26%±0.9 90 minutes post CNO injection; **Figure 4d, e and f; Supplementary Fig. 4a, b and c**).

In conclusion activation of C-LTMRs transiently increases all kind of social interaction between animals to the expense of isolated behaviour for up to 90 minutes post CNO injections, including behaviour related to skin to skin contacts and social exploration. The impact of such behavioural alteration appears to impact group dynamics for longer periods, especially groups of 3 mice. Remarkably, some behavioural traits were still significantly increased up to 20 hours after CNO injection, such as the duration of nose-to-nose interaction, social approach and stops in contact (**Figure 4e**) as well as the number of social approach and social escape (**Figure 4d**). In contrast, all behavioural traits in C-LMTRs^hM3Dq^ mice came back to control mice level after 20 hours post injection.

Next, the relationship between each mouse and the group dynamics was analysed. First, we focused on group of two mice and, in our condition, one mouse of a given genotype had a probability of 1/3 to interact with a mouse from the same genotype, and 2/3 to interact with a mouse from the other genotype (**Figure 5a**). Remarkably, after CNO but not saline injection, C-LMTRs^hM3Dq^ mice interacted more with each other than expected by chance level (+11.7%±0.2 compared to saline), while Control mice interacted less with each other (- 9.8%±0.2 compared to saline) (**Figure 5b and c**). In addition, Control mice had a higher probability of contacting C-LMTRs^hM3Dq^ mice than expected (+9.7%±0.3 compared to saline). This effect was visible for 30- and 90-minutes post CNO injection, depending on the couple formed. These observations are only valid when all types of contacts are analysed as a whole but not for individual behaviours related to social exploration where only C-LTMRs^hm3Dq^ mice had a higher probability of interacting with each other (**Supplementary Fig. 5**). The mean duration of the time spent in a group of 2 was however similar. Next, we focused our investigation of the dynamic of groups of three mice. While looking at the combination of mouse making or breaking groups-of-three, we did not observe any differences after CNO injection compared to saline (**Figure 5d and f**), and this held true at all time point. However, it appears that the groups of three mice formed by a C-LMTRs^hM3Dq^ mouse lasted longer than those created by a Control mouse, especially if the C-LMTRs^hM3Dq^ mouse joined two other animals to create a mix group (one C-LMTRs^hM3Dq^ mouse and two Controls) (**Figure 5e**_**2**_). This effect was particularly striking 1 hour after CNO injection, but decreased after 90 minutes and was completely abolished after 390 minutes (**Figure 5e**_**3**_).

**Figure 5:**
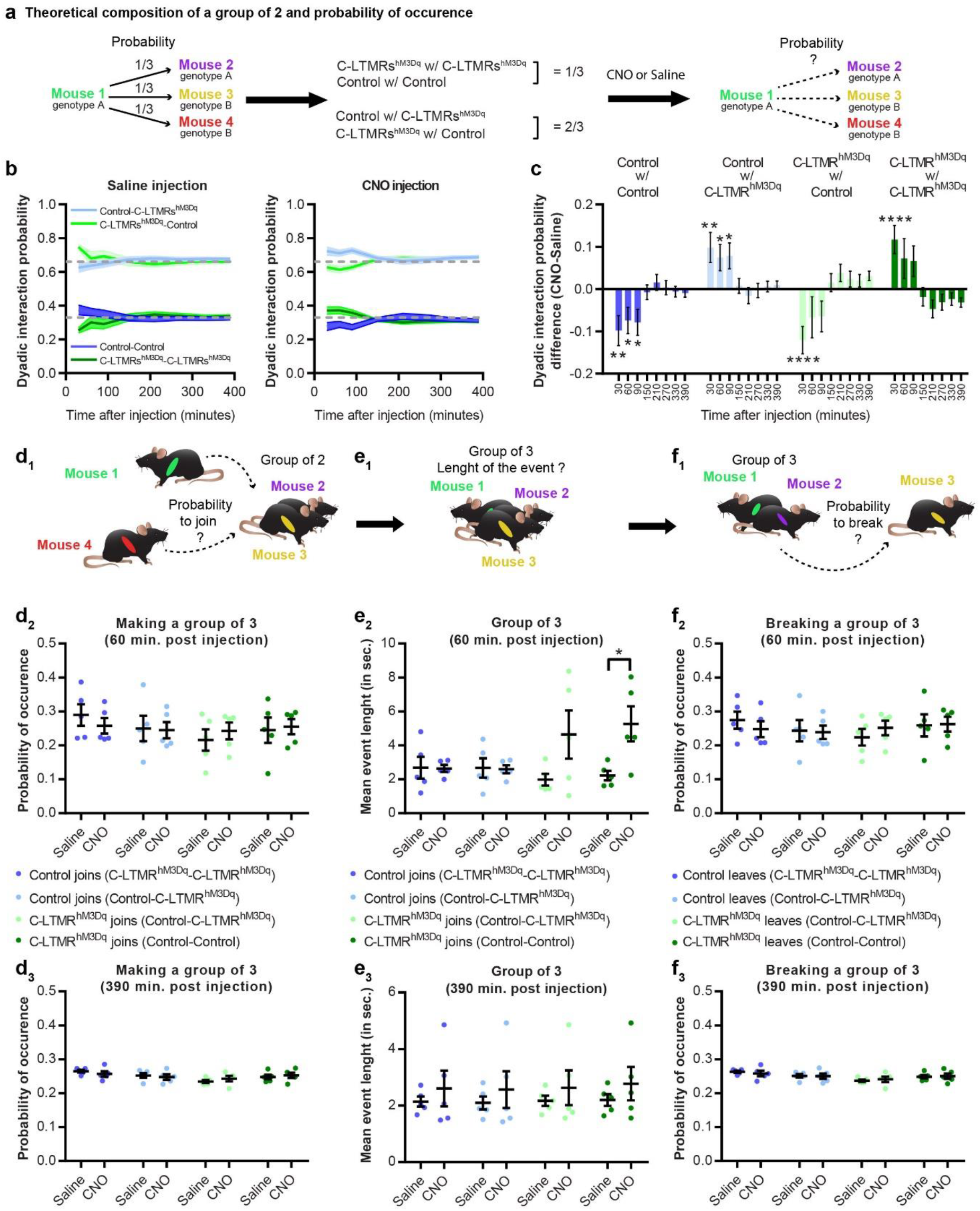
C-LTMRs exogenous activation shakes group dynamics and inter individual interaction probabilities. a. Theoretical composition of a group of two based on its probability of occurrence b. Dyadic interaction probability for all 4 possibilities at 30, 60, 90, 150, 210, 270, 330, and 390 minutes after saline (left) and CNO (right) injection. a. c. Dyadic interaction probability difference between CNO and saline injections at 30, 60, 90, 150, 210, 270, 330, and 390 minutes post injection. two-way RM ANOVA, Bonferroni post hoc test, ∗p < 0.05; ∗∗ p<0.01; ∗∗∗∗p<0.0001 d. Probability of occurrence for a mouse of any given genotype to create a group of three of all possible composition at 60 (d2) or 390 (d3) minutes post saline or CNO injection. e. length of the group 3 event created by a given mouse depending of the total composition of the group at 60 (e2 or 390 (e3) minutes post saline or CNO injection. f. Probability of occurrence for a mouse of any given genotype to break a group of three of all possible composition at 60 (f2) or 390 (f3) minutes post saline or CNO injection. Paired t test p= 0.0486

## Discussion

Understanding how touch shapes social interactions, while keeping a level of ethological validity is particularly challenging. In this study we overcame this challenge by combining a unique genetic strategy and new tracking technologies to characterize the contribution of C-LTMRs to affective and social touch in mice. By reducing, just before birth, the activity of this specific population of primary sensory neurons defined genetically and physiologically, we observed in adults a reduction of contacts with other mice, leading to an increase in isolated behaviours. Conversely, exacerbation of C-LTMR excitability with a chemogenetic approach in adults, led to an increase in social interaction, reduced isolated behaviour, and changed groups social dynamics. We present evidence that C-LTMRs may be one of the main contributors to social development and a potential target of treatment for neurodevelopmental disorders.

### C-LTMRs, Ca_v_3.2 and sociability

Even if the C-LTMR hypofunction and the C-LTMRs remote activation share a similar genetic strategy, the behavioural phenotypes observed in adults may results from totally different mechanism. C-LTMRs inhibition using Ca_v_3.2 cKO^Nav1.8^ mice is a conditional knock-out where pro-excitatory calcium channel Ca_v_3.2 expression is removed in C-LTMRs as soon as Nav1.8 promoter and the Cre recombinase start to be active, around E16-E17 ^29,30^. Thus, this “late” DRG Cav3.2 conditional KO spare any in utero development presumably dependent of Cav3.2, while it starts to abolish Cav3.2 function in neuronal excitability at perinatal stage. We and other documented the impact of Ca_v_3.2 in helping LTMR neurons to fire in burst by lowering action potential threshold and by generating an after depolarisation potential ^16,31–33^. Accordingly, C-LTMR remained mechanosensitive after Ca_v_3.2 conditional knock out, but responded to higher threshold stimuli as the result of dampened excitability ^16^. Touch perception defects from birth may have dramatic consequences on nurturing touch which can lead to impaired somatosensory development, increased stereotyped behaviours and deficits in social behaviour and cognitive abilities ^34–36^. Because of the affective nature of the information carried by CTs, and potentially their rodent equivalent, C-LTMRs, it has been hypothesized that these neurons play a critical role in this process. Interestingly, Ca_v_3.2^Nav1.8^cKO mice share some phenotypes with two mouse models of ASD which have been phenotyped with the LMT (Shank 2 and Shank 3 KOs)^21^. Specifically, Shank 2 KO and Ca_v_3.2^Nav1.8^cKO mice share common phenotypes with regards to isolated behaviours and the contact reduction. On the other hand, C-LTMRs Ca_v_3.2 cKO mice do not show any deficit in exploratory behaviours **Figure 1 c and d and Supplementary Fig. 1**)^21^. The fact that no difference was observed in either social escape behaviours or in passive social interaction in these mice while unilateral and reciprocal interactions were decreased also suggests that Ca_v_3.2^Nav1.8^cKO mice do not seek to avoid inter-individual tactile stimuli, but rather do not actively seek it. Nonetheless, if these mice do not show signs of tactile avoidance, based on the parameters automatically extracted by the LMT and on the Crawley test results where only the time of interaction is affected, this could be due to the experimental paradigm. Indeed, it may be difficult for a mouse to avoid 3 others in the LMT arena (50∗50cm).

It is also interesting to note that Ca_v_3.2^Nav1.8^cKO mice have an excessive rearing behaviour when isolated (**Figure 1 c and d**), which can be considered a manifestation of anxiety ^37^. Rearing behaviour’s relationship with anxiety level is a matter of debate, this phenotype, associated with decreased sociability in the LMT and in the Crawley test is strikingly similar to those observed in the Mecp2 and Shank3 peripherally restricted KO ^14^. These observations strongly suggest that C-LTMRs are a key component of social interactions and may play a critical role in neurodevelopmental disorder such as ASD, as suggested from human observation. Indeed, individuals with ASD have altered tactile sensitivities and autism-associated behavioural deficits and neural responses to C-LTMR-triggered affective touch stimuli are inversely correlated ^38^. Such observations suggest that people with greater numbers of autism-relevant traits have impaired processing of affective touch. Because most young children with ASD are averse to touch, caregivers often provide less nurturing touch, and this lack of tactile input may have a profound impact on subsequent behaviour and development^39^.

The asocial phenotypes observed in this study in adult C-LTMRs Ca_v_3.2^Nav1.8^cKO could then be the reflect of long-term consequences of altered bottom up effect of touch on shaping the developing social brain during early life. Finally, the asocial traits revealed here may have translational relevance in clinic. Indeed, they nicely parallel the presence of congenital missense mutations in distinct ASD patients within the Cacna1H gene leading to Cav3.2 functional defects ^40^. While in this case, mutations have body wide consequences, we can speculate a substantial contribution of tactile sensory deficits to explain the genotype-phenotype relations. Consequently, a perspective of developing peripherally restricted Cav3.2 selective T-type calcium channel activators could represent a therapeutic opportunity to correct early ASD defects. Conversely, caution should be warranted with the use of T-type calcium channels inhibitors specifically with the risk of hitting the critical period of social brain development in child’s early life.

### C-LTMRs and pleasant touch

Using a model of chemogenetic activation of C-LTMRs we demonstrated that this population of neurons is sufficient to create a pleasant experience. Not only does C-LTMR stimulation induce a rewarding experience in a CCP paradigm with no specific context, but it also reinforces thermotaxis centred around 30°C, consistent with the idea that the functioning of these neurons is tuned to the temperature of a skin-stroking caress (∼30°C) as in humans ^41^. Consistently, we previously documented that Cav3.2^Nav1.8^cKO mice has the exact opposite phenotype with a weakened thermotaxis ^16^. Furthermore, the temperature at which the LMT experiments are performed is 24°C, which is cool for mice. Therefore, the CNO reinforcement of thermotaxia evidenced in the gradient paradigm, is likely to contribute to the animal seeking for group formation within the LMT where the warm body temperature of each other mouse in duo, trio, or even tetrad further amplify the C-LTMR activation during skin to skin contacts. This data reinforces our confidence that the population we defined as C-LTMRs by using different gene expression such as TAFA4, TH and the duo Ca_v_3.2-Na_v_1.8 is the correlate of the sensory fibres supporting affective touch in humans.

It is interesting to note that the mini Ca_v_3.2 promoter used in our viral constructs may not be as efficient as those usually used in similar strategy such as CAG, hSyn, or Ef1A. By contrast, our approach likely results in hM3Dq expression levels just sufficient to potentiate C-LTMRs excitability rather than over-excite them, thus explaining the subtle behaviour changes observed. In vitro calcium imaging in cultured DRG from DREADD expressing mice confirms that most of the neurons functionally responding to DREADD agonist CNO had a unique C-LTMR chemo response pattern consistent with their transcriptional profile regarding the expression of channels and receptors coupled to cytosolic calcium variations. This includes responses to agonists of TRPA1 agonist as reported before ^15,16^, as well as to the purinergic metabotropic receptor P2RY1 that is consistently reported to be highly expressed in C-LTMRs across species from rodents to non-human primates ^18,20,42,43^. In addition, the CNO responsive neurons where all negative to live staining with fluorescent IB4, as expected from previous studies ^15^.

To our knowledge, only one study presently published has been able to demonstrate that a population of primary sensory neurons, the population expressing MRGPRB4, is able to drive a motivational behaviour in mice upon DREADD stimulation approach ^44^. Unfortunately, whether or not this neuronal population can also drive specific inter-individual behaviours has not been investigated.

Whether the genetically defined C-LTMR population MRGPRB4 or TH/VGlut3/TAFA4 (or both) is the murine equivalent of C-tactile fibres in humans remains controversial. MRGPRB4-expressing population is strikingly different from the TH-expressing C-LMTRs population. Despite both belonging to a class of mechanoreceptive C-fibres, MRGPRB4 is expressed in a population not as well defined functionally and genetically across development than TH-expressing neurons ^45,46^. For example MRGPRB4-expressing neurons also express TRPV1 and respond to capsaicin ^44^ similarly to numerous C-nociceptors, whereas TH-expressing neurons do not ^20^. Interestingly humans C-Tactile fibres do not seems to be sensitized by capsaicin, suggesting that they do not express TRPV1 ^3^. In addition, T-type channel blocker TTA-A2 applied in human glabrous skin completely abolished sensibility to innocuous tactile stimuli supporting expression of T-Type calcium channel in C-Tactile afferences, similar to mouse TH-expressing C-LTMRs, but not MRGPRD4 fibres ^16,47^.

### C-LTMRs as a motivational drive toward social interaction

Activation of C-LTMRs greatly increase the number of contacts between adult animals. In accordance with C-LTMRs known innervation of the hairy skin, we observed a large increase of what we considered skin-to-skin contact (contact *side by side* and *side by side opposite*, cumulative duration and number of events **Figure 5 d-e**)^17^. More surprisingly, we also witnessed an increase in nose-to-nose contacts and even more nose-to-anogenital contacts, involving skin areas that are not supposedly innervated by C-LTMRs, although recent work suggest that C-LTMRs innervates more skin areas than expected in humans, such as glabrous skin ^48^. Such an observation may be due to a priming or reinforcing influence of C-LTMRs activation on inter individual interaction to induce more complex social contacts such as social investigation (nose to nose and nose to anogenital contacts). For example, C-LTMRs stimulation via side to side contact, may act as an appeasing signal to engage communication through face to face sniffing or may be the starter for soliciting play behaviour such as pounce and crossover (not annotated by the LMT)^49,50^. This may also explain the surprising results concerning the changes observed in dyadic and triadic interactions dynamics where C-LTMRs activation in one mouse reinforce social interaction with the other C-LTMR-stimulated mouse as well as with control mice where C-LTMRs are not potentiated. The LMT is a close arena where all four mice are free to interact with each other. Consequently, a change in occurrence of one given social behavioural trait in any mouse logically impact the rest of the mice, including control mice. Thus, explaining the observed significative alteration of interaction probability of Control mice with C-LTMRs. In addition, C-LTMR-stimulated mice may act as social stress buffers, appeasing social tension in the whole group, thus increasing seeking for inter-individual contact in all animals, including in Controls. However, our results suggest that even if Control mice behaviour appeared to be impacted by the CNO treatment of all four mice, the effect is more robust and potent in C-LTMR-stimulated mouse, suggesting that a C-LMTRs^hM3Dq^ mouse will have a preference toward another C-LMTRs^hM3Dq^ mouse over control mice after CNO treatment. Thus, reciprocal stimulation of C-LTMRs may be more reinforcing than unilateral stimulation.

In conclusion, while clinical studies documented that affective information conveyed through the skin have powerful impact on social behaviour, the direct causality of C-Tactile/C-LTMRs primary afferents in this bottom up social brain regulation remained to be rationally demonstrated. Our study, combining mouse genetics and in-depth ethological analysis, provides a first demonstration that in healthy naÏve adults, enhancing skin C-LTMRs activity for a few tens of minutes is sufficient to induce immediate and lasting prosocial effects, while conversely impairment of C-LTMRs functions from birth to adulthood negatively impact sociability with behavioural traits resembling those found in genetic mouse models of Autistic Spectrum Disorders. We hope that the type of preclinical approaches developed here to modulate C-LTMRS with a level a selectivity that has not been yet achieved, will prefigure future investigations aimed at better understanding the pathophysiology of CNS circuits of social behaviours driven by affective touch.

## Acknowledgments

We would like to thank the PVM viral vectorology facility of Montpellier, and the Canadian Neurophotonics Platform Viral Vector Core Facility for the production of viruses, John Wood for the Nav1.8-Cre mouse line, Jean Chemin for transient transfection of HEK293T cells, Emmanuel Valjent for sharing data in DRGs from the Ribotag-HA reporter mice, Luc Forichon, Steeve Thirard and the IExplore-RAM animal facility for their crucial help, Emmanuel Valjent for his help with the CPP, Reda El Mazouz for its help in calcium imaging study, Aziz Moqrich for continuous support and advices on the study of C-LTMRs, Fabrice de Chaumont for advices on the LMT and social behaviour, Muriel Asari and SciDraw.Io (doi.org/10.5281/zenodo.3925997)for the illustrations, Etienne Audinat and Yan Emery for reading the manuscript. This work has been supported by the Agence National pour la Recherche (grants ANR15-CE-16-012-Pain-T and Labex ICST to EB), the Fondation pour la Recherche Médicale (équipe FRM Pain-T grant to EB), the International Association for the study of Pain (early career research grant to AF), the Centre national de la recherche scientifique (CNRS), l’Institut national de la santé et de la recherche médicale (INSERM), and the University of Montpellier.

## Authors contribution

AF conceptualized performed and analysed most of the experiments. EB and PF helped to conceptualize the experiments and designed the AAV vectors. DH and FJ helped with the LMT experiments. EB performed the calcium imaging on dissociated DRG. MM performed and analysed the three-chamber social interaction test. AF wrote the manuscript with the help of all the authors.

## Methods

### Study approval

All animal procedures complied with the welfare guidelines of the European Community and were approved by the local ethic comity, the Herault department Veterinary Direction, France and the French ministry for higher education, research and innovation (Agreement Number:2017100915448101).

### Animals

Ca_v_3.2 KI ^GFP-flox^, Cav3.2^Nav1.8^cKO (Cav3.2 KI ^GFP-flox^ x Nav1.8^cre^) and Nav1.8^cre^ mouse lines were bred and housed in a Specific Pathogen Free (SPF) animal facility at the Institute of Functional genomic under approved laboratory conditions (12 hours day/night cycle. 22–24 °C, 50 ± 5% humidity, food and water *ad libidum*). All behavioural tests were conducted in the same animal facility with the SPF sanitary status.

### Plasmids and Viruses

The excitatory Dreadd hM3Dq with a N-terminal HA tag epitope was cloned into a pAAV viral vector under the control of a 1.5kb minimal Cav3.2 promotor ^23^ and between double-floxed inverse Orientation (also called FLEx) sequences, allowing to switch the Dreadd open reading frame in the correct orientation upon Cre mediated recombination. The pAAV-promCav3.2-DIO-ChR2-ires-YFP-WPRE ^23^ served as template. The ChR2-ires-YFP was replaced by the HA-hM3Dq sequence, coming from pAAV-hDlx-GqDREADD-dTomato (gift from Gordon Fishell, Addgene plasmid # 83897), using HIFI DNA assembly method (New England Biolab, Evry - France). A 3xHA version of the Dreadd was further created by inserting a KpnI-NheI cassette obtained by gene synthesis (Proteogenix, Strasbourg - France). All constructs were verified by DNA sequencing. Before viral production, the functionality of the constructs was demonstrated by Fura2 calcium imaging following transient transfection in HEK cells together with expression vectors expressing the Cre recombinase (pCAG-Cre-ires-GFP, a gift from Jérôme Dujardin), and the Egr1 transcription factor (pcDNA3-Egr1, a gift from Eileen Adamson, Addgene plasmid #11729) known to stimulate the Cav3.2 promotor ^25^.

AAV-PHPs viruses were custom made either by the Plateforme de Vectorologie de Montpelier (PVM, Montpellier, France) or by the Canadian Neurophotonics Platform Viral Vector Core Facility (RRID:SCR_016477) (Quebec - Canada) using the ad hoc capside plasmid (pUCmini-iCAP-PHP.S, a gifts from Viviana Gradinaru; Addgene plasmid # 103006; ^24^). Control rAAV PHPs CAG mCherry was bought from Addgene (viral prep # 59462-PHP.S)

### Primary sensory neuron culture and calcium imaging

Lumbar/thoracic DRGs (L4 to T10) were prepared and cultured on Laminin coated µ-Dish chambers (Ibidi, Germany) as described previously (Francois et al., 2013) from AAV injected mice (18 weeks) Recording were made within 24h of culture. Prior to recording, neurons were incubated with 5μM fura-2AM in Tyrode’s solution for 1 hour at 37°C, then washed and kept for an additional 20 minutes in Tyrode with no Fura2 for desesterification. Fluorescence measurements were sampled at 1Hz with an inverted microscope (Olympus IX70) equipped with an Evolves Photometrics EMCCD camera (Roper Scientific, France). Fura-2 was excited at 340nm and 380nm and ratios of emitted fluorescence at 510nm were acquired using Metafluor software (Universal Imaging). Drugs were applied with a gravity driven perfusion (1-2ml/min) Pharmacological agonists of hM3Dq (CNO 30µM, Tocris, France), P2YR1 (MRS2365 200nM, Tocris France), TRPV1 (Capsaicin 500nM), TRPA1 (allyl isothiocyanate 200µM), KCl (40mM) were prepared into the Tyrode solution and applied sequentially to the neurons for a few seconds. Data were analyzed offline using metafluor, excel, and graphpad. Similar procedure for Fura2 loading and imaging was used for transiently transfected HEK cells for the verification of pAAV functionality prior to custom virus production. Otherwise stated, all chemicals were from Sigma Aldrich (L’isle d’Abeau Chesnes - France).

### Intrathecal injection and Stereotaxic surgeries

6 to 8week old mice were anesthetized by inhalation of a 2%isoflurane/1.5% oxygen mixture.

For intrathecal injection, 5µl of viral vector (1.10^13^) was injected into the lumbar subarachnoid space using a 20µl Hamilton syringe and 26g removable minimal dead volume needle. Wounds were sutured and surgical sites infiltrated with 2% lidocaine in saline. Animals were placed on a heating pad in an oxygenated chamber and monitored until fully recovered.

### Behaviour

#### Three chambers social interaction test

8 to 12 weeks old male mice were first habituated for half an hour to the experimental room in their home cage. The three-chamber arena is composed of 3 compartment box, each 20×40×22(h)cm and separated by two sliding doors (5×8(h)cm) with one prison placed in the upper right corner of the right compartment and another one on the lower left corner of the left compartment. After room habituation, each animal was placed individually in the arena, starting from the middle compartment and was let free to explore for 10 minutes with the two prisons empty. Then, the mouse was locked in the middle compartment and one mouse, stranger to the tested mouse (C57B6J/n bred and housed in the IGF facility), was place in one of the two prison and an inanimate object (made from Lego blocks, roughly shaped like a mouse) was placed on the other prison. The tested mouse was then set free to explore all compartments again for 10 minutes. The all experiment was video recorded and animal position was tracked using Ethovison XT13 (Nodlus). The preference index was calculated with the following formula (Time in stranger side - Time in object side)/exploration time. A mouse was considered interacting automatically by Ethovision when the tested animal was facing the middle of the prison inside a 1cm perimeter around the prisons. The side where the stranger mouse and the inanimate object were placed was alternated between each tested mouse to avoid bias. Experiments were performed in the morning (from 7 to 10 a.m.) under 180 lumens. The experimenter was blinded to the animal conditions.

#### Live Mouse Tracker

The LMT setup was built following the instruction of De Chaumont et al., ^21^. We added to the original design two water dispensers to the arena on two sides.

Before going into the LMT each group of 4 mice were housed together for at least 2 weeks (following RFID chip implantation under anaesthesia). LMT recordings were performed with 10 to 16 weeks old Ca_v_3.2^GFP-flox^KI or Cav3.2^GFP-flox^KI x Nav1.8^cre^ mice, and with 14 to 18 weeks old Nav1.8^cre^ mice (injected with rAAV_PHPs_-CAG-mCherry or rAAV_PHPs_-Ca_v_3.2-FLEx-HA-hM3Dq). During recording, the animals were kept under the same condition as in the housing facility (12hour Daylight, 500 lumens, food and water *ad libidum*) and the experimenter came once per day to perform injection if necessary or to check the water and food level.

We used Python scripts provided to analyse all the data acquired by the system. (www.livemousetracker.com; https://github.com/fdechaumont/lmt-analysis). We only performed analyse during the activity phase (night cycle) as during the day mice nest together which impaired the tracking and provide unreliable annotation.

#### von Frey

8 weeks after intrathecal injections, mice were first habituated for half an hour to the experimental room in their home cage. Each mouse was then placed individually into small arena (8∗8cm) over a von Frey mesh for 45 minutes for habituation. Then, animals were injected peritoneallly with 1mg/kg CNO diluted in sterile saline (prepared fresh each time). 30 to 45 minutes after the injections, von Frey filaments (0.07g, 0.6g, 2g) and the brush was applied 5 time on each hind paw and withdrawal was then scored. Experiments were performed in the morning (from 8 to 11 a.m.) under 500 lumens. The experimenter was blinded to the animal conditions.

#### Gradient

8 weeks after intrathecal injections, mice were first habituated for half an hour to the experimental room in their home cage and then injected with 1mg/kg CNO diluted in sterile saline (prepared fresh each time). 30 minutes later, mice were placed into the bioseb thermal gradient (2 corridors of 1m50 with one extremity cooled down to 5°C and the other to 50°C, creating a thermal gradient spited into 20 thermal zone). Mice were free to explore for 90 minutes during which animal’s positions were tracked and annotated accordingly to the temperature zones. For each run, a C-LTMRs^hM3Dq^ mouse and a control mouse were tested simultaneously. However, the groups were labelled as such as he experimenter was still blind to the animal conditions. Mice corridor distribution was alternated between each run to avoid any corridor biased. Experiments were performed in the morning (from 8 to 12 a.m.) under 180 lumens.

#### Conditioned place preference (CPP)

8 weeks after intrathecal injections, mice were subjected to the conditioned place preference protocol. This protocol consisted into 6 experimental days: the first day consisted into an exploration and habituation phase of 30 minutes per mouse where mice were free to explore the CPP arena (compartments: 25cm∗20cm, corridor: 5cm∗20cm . From the second to the fifth day, animals were conditioned to associate a saline injection in the morning to their preferred compartment (defined during the first day), and a CNO injection (1mg/kg) to the other comportment in the afternoon. During this conditioning, animals received the injection and stayed in their homecage for 30 minutes and then were placed into the designated compartment for 30 minutes, to allow the CNO to reach its molecular target. To avoid that the CNO was still active between two conditioning, we choose to stretch as much as possible the period of time between the CNO injection from one given day and the next day saline injection. On the sixth and last day animals were free to explore the all arena again for 30 minutes. The position of the animal was detected by a network of infrared beams. The CPP rack allowed us to perform 4 experiments at a time (Imetronic place preference setup), thus each run was balanced to test 2 C-LTMRs^hM3Dq^ and 2 control mice, however the groups were labelled as such as the experimenter was blinded to the animal conditions. Experiments were performed in the morning (from 8 to 12 a.m.) under 180 lumens.

### Immunohistology

Tissue collection and processing. Animals were transcardially perfused with phosphate-buffered saline (PBS) followed by 10% formaldehyde in PBS. Brain and spinal cord were dissected, post-fixed in 10% formaldehyde for 24 hours, and cryoprotected in 30% sucrose in PBS. DRGs were cryoprotected directly after formaldehyde perfusion. Tissues were then frozen in Optimum Cutting Temperature (OCT, Tissue Tek) and sectioned using a cryostat (Leica). Spinal cord and brain were sectioned at 40μm and stored in PBS + 0.05% azide at 4°C. For DRGs, tissues were sectioned at 18 μm, collected on Superfrost Plus slides (Fisher Scientific), and stored at −80°C.

Immunofluorescence. Tissues were incubated for 1 hour and blocked in a solution consisting of 0.1 M PBS with 0.3% Triton X-100 (Sigma) plus 5% normal donkey serum. Primary and secondary antibodies were diluted in 0.1 M PBS with 0.3% Triton X-100 plus 1% normal donkey serum. Sections were then incubated overnight at 4°C in primary antibody solution, washed in 0.1 M PBS with 0.3% Triton X-100 for 40 min, incubated for 2 hrs in secondary antibody at room temperature (RT), and washed again in 0.1 M PB for 40 min. Sections were then mounted using Dako fluorescence mounting medium. Images were acquired with a Leica SP8 confocal microscope.

Primary antibodies: anti-TH: Millipore (sheep; 1:500), anti-HAtag: Covenant (mouse, 1:1000); Anti-Vglut3 : Synaptic system (Rabbit, 1:500). To identify IB4-binding cells, Fluorophore-conjugated IB4 (VectorLab, 1:500) was used in place of primary and secondary antibodies. Secondary antibodies: Alexa Fluor®-conjugated secondary antibodies were acquired from Invitrogen and Jackson Immunoresearch Labs.

### Statistics

Statistics were performed with Graphpad Prism 8. Data are represented as mean ± s.e.m. For bar graphs, each individual data points were superimposed under mean mean ± s.e.m. Statistical tests used to compare values are indicated in each figure legends.

---

**Supplementary Figure 1:**
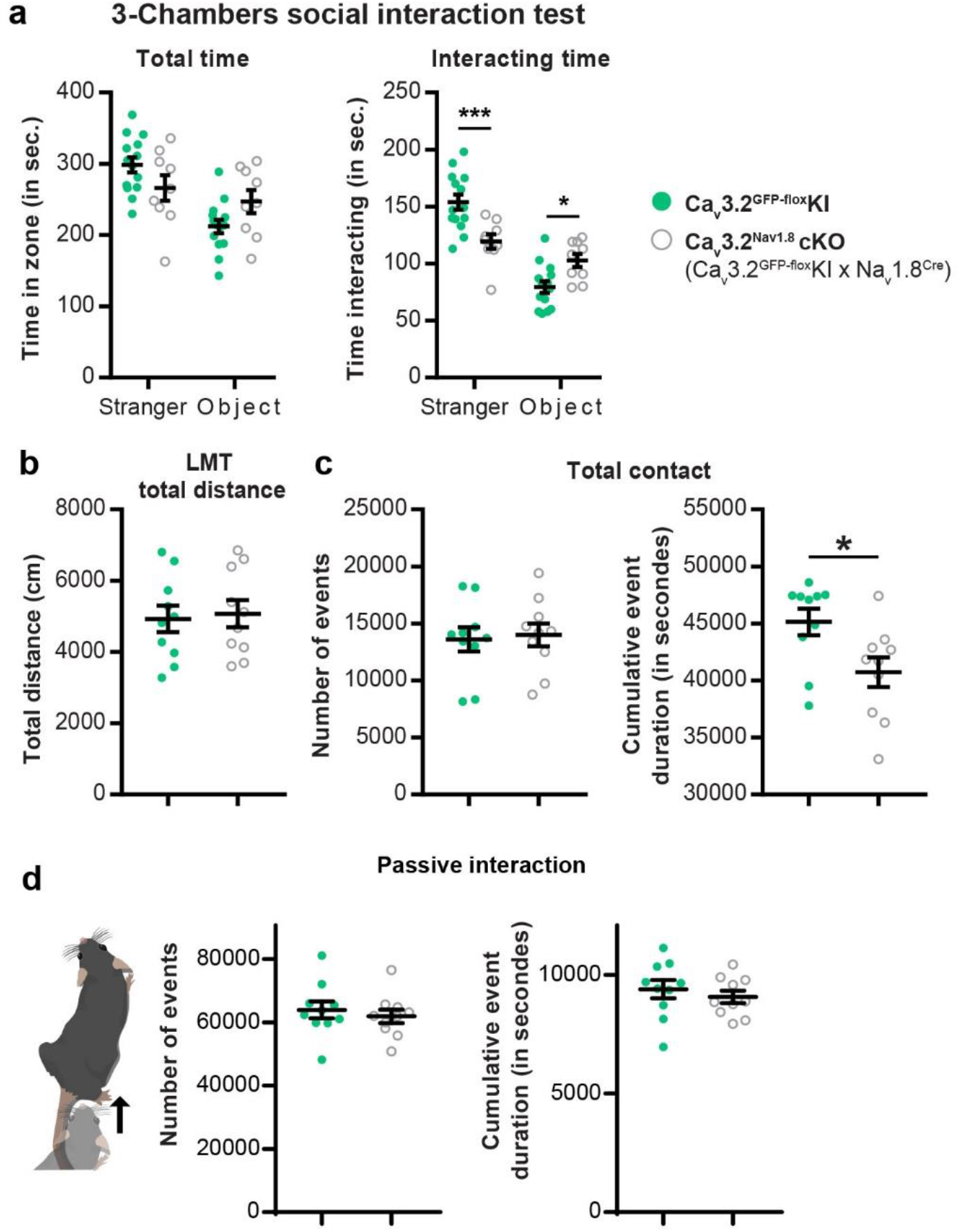
C-LTMRs hypo excitability via Ca_v_3.2 deletion reduces social behaviour. a. In a three-chamber social interaction test, control Ca_v_3.2^GFP-flox^KI littermates spent more time interacting with a stranger mouse and less time with an inanimate object than Ca_v_3.2^Nav1.8^cKO. 2 way ANOVA with Bonferroni post hoc test, ∗∗∗p=0.0007; ∗ p=0.262, n Cav3.2 KI ^GFP-flox^ = 14 (light green circles), n Cav3.2 KI ^GFP-flox^ = 9 (open grey circles). b. Sum of the total distance travelled by each mouse during the three nights for Ca_v_3.2 KI ^GFP-flox^ KI (light green circles; n =10) and Ca_v_3.2^Nav1.8^cKO (open grey circles; n=10). c. raw values extracted from the LMT for three consecutive nights. Left: number of total contact, right, cumulative event duration of contacts (∗ p=0.0220). d. number of events and cumulative duration of passive social interaction. Wilcoxon matched-unpaired t-test

**Supplementary Figure 2:**
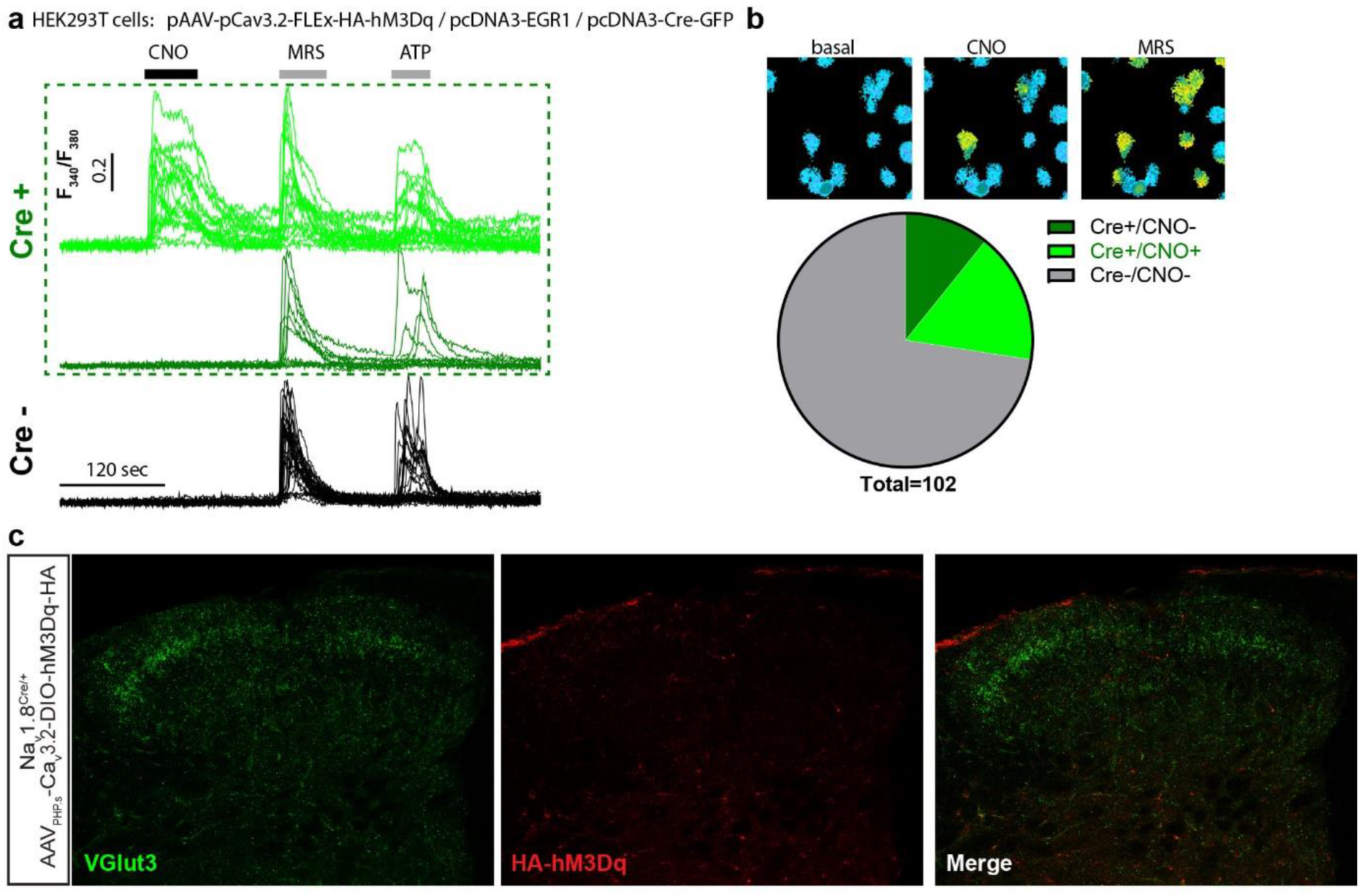
validation of the viral construct used to target C-LTMRs. a. Representative individual Fura2 calcium influx in cultured tsA201 cells transfected with pAAV-pCav3.2-FLEx-HA-hM3Dq, pcDNA3-EGR1, and pcDNA3-Cre-GFP, following bath perfusion of CNO (30µM), MRS2365 (200nM) to stimulate endogenous P2Y1 receptors or ATP (30µM) to stimulate all endogenous P2Y receptors. Green traces: cells expressing the Cre recombinase (visualized with the GFP). Black traces: cells that do not express the Cre recombinase. b. Top panels, representative images of ratiometric calcium imaging on tsA201 cells transfected with pAAV-Cav3.2-FLEx-HA-hM3Dq, pcDNA3-EGR1, and pcDNA3-Cre-GFP following bath perfusion of CNO to stimulate hM3Dq (30µM, middle), and MRS2365 (200nM, right). Bottom panel: Pie chart of the number or cells responding to CNO perfusion. 102 cells recorded c. Representative images of VGlut3 (Green) and HA (Red) immunostaining in the caudal part of the lumbar spinal cord.

**Supplementary Figure 3:**
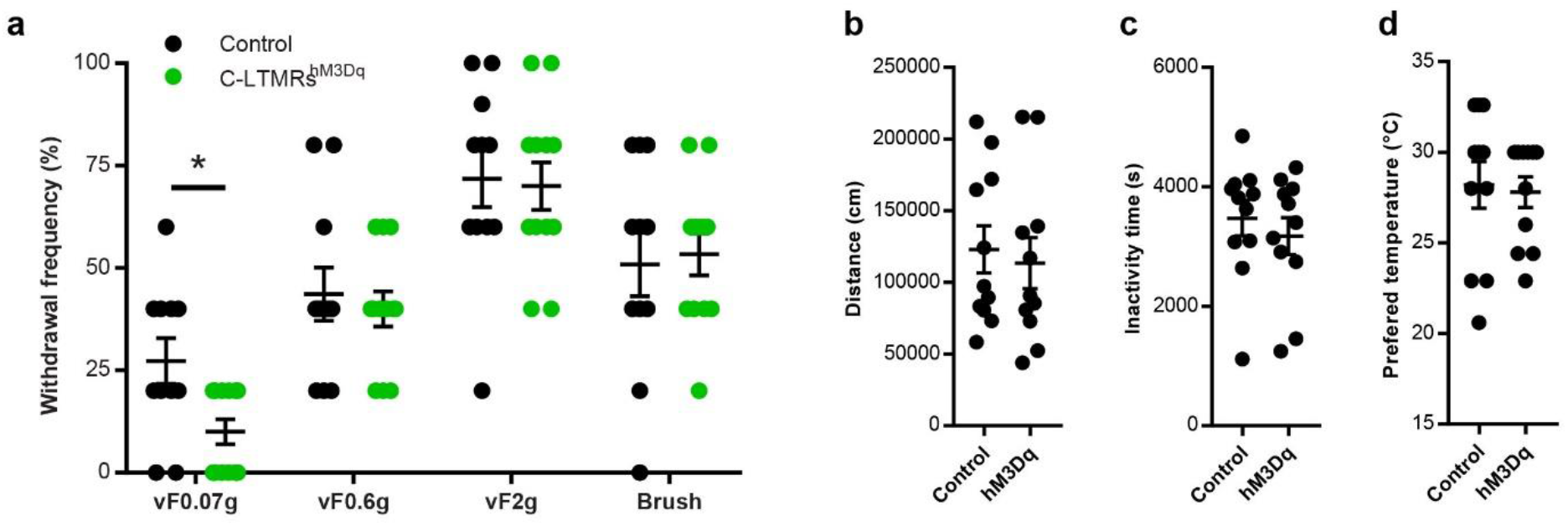
Light touch perception is altered by C-LTMRs exogenous activation. a. Withdrawal frequency to von Frey filament 0.07g, 0.6g and 2g of Control (Nav1.8^Cre^; AAV_PHPs_-CAG-mCherry; in black) and C-LTMRs^hM3Dq^ (Nav1.8^Cre^; AAV_PHPs_-pCa_v_3.2-FLEx-HA-hM3Dq; in green) mice. Mann-Whitney test; ∗ p=0.0221; Control n =11, C-LTMRs^hM3Dq^ n =12) b. Distance travelled during 90 minutes on the thermal gradient. c. Cumulative time of inactivity during 90 minutes on the thermal gradient. d. Average of the temperature where the animals spent the most of their time.

**Supplementary Figure 4:**
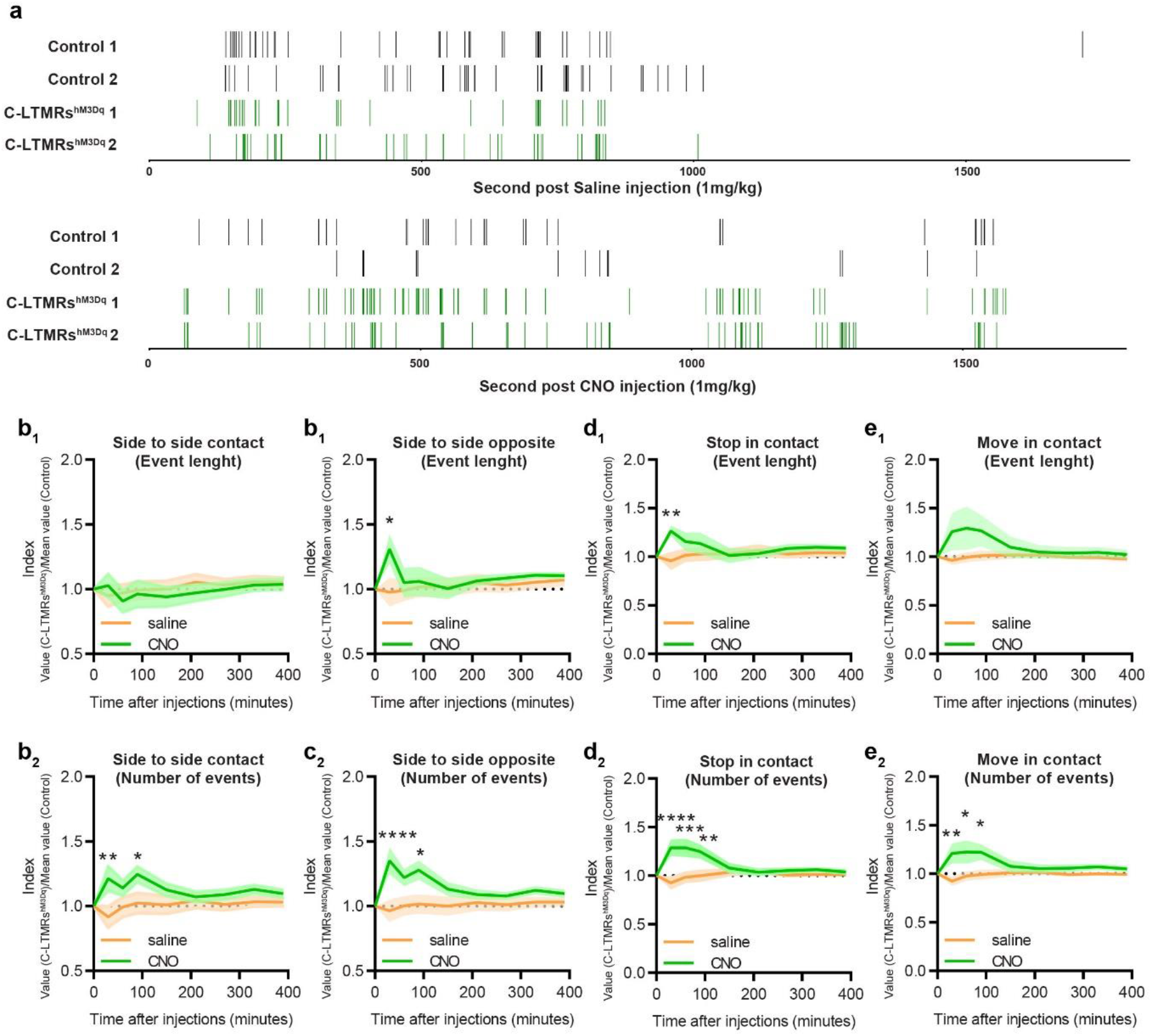
C-LTMRs exogenous activation transiently increases social exploratory contacts between animals. a. Example from the 4^th^ group of nose-to-nose contacts events for the first 30 minutes post saline (upper panel) or CNO (lower panel) injections. b to e. LMT index obtained at 30, 60, 90, 150, 210, 270, 330, and 390 post saline injection (orange) or post CNO injection (green) for the cumulative number of contact side to side (b2) and their cumulative length (b1), the cumulative number of contact side to side opposite (c2) and their cumulative length (c1), the cumulative number of events stop in contact (d2) and their cumulative length (d1), or the cumulative number of event move in contact (e2) and their cumulative length (e1). two-way RM ANOVA, Bonferroni post hoc test, ∗p < 0.05 ; ∗∗ p<0.01; ∗∗∗ p<0.005; ∗∗∗∗p<0.0001

**Supplementary Figure 5:**
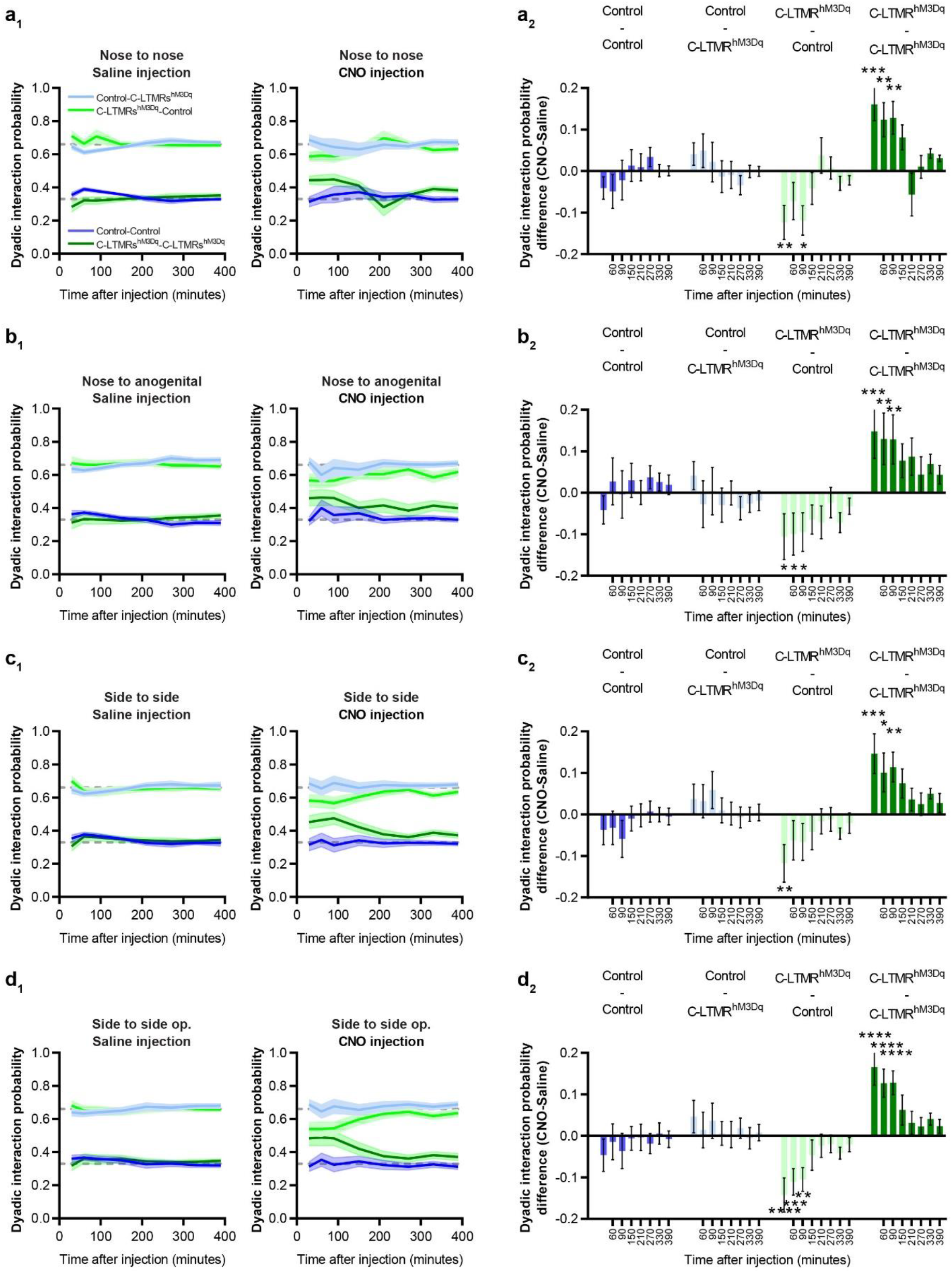
C-LTMRs exogenous activation exacerbates interaction probabilities between C-LTMR^hM3Dq^ mice. a1 to d1. Specific dyadic interaction probability for all 4 possibilities at 30, 60, 90, 150, 210, 270, 330, and 390 minutes after saline (left) and CNO (right) injection. A1 nose to nose dyadic interaction; b1 nose to anogenital dyadic interaction; c1 side to side dyadic interaction; d1 side to side opposite dyadic interaction. a2 to d2. Specific dyadic interaction probability difference between CNO and saline injections at 30, 60, 90, 150, 210, 270, 330, and 390 minutes post injection. a2 nose to nose dyadic interaction; b2 nose to anogenital dyadic interaction; c2 side to side dyadic interaction; d2 side to side opposite dyadic interaction. two-way RM ANOVA, Bonferroni post hoc test, ∗p < 0.05 ; ∗∗ p<0.01; ∗∗∗∗p<0.0001

